# Characterization of household microbiomes from three unique cities around the world

**DOI:** 10.64898/2026.04.11.717928

**Authors:** Caroline Scranton, Victoria Obergh, Madison Goforth, Kavitha Ravi, Poornima Jayakrishna, S.K. Anisa, Stephanie A. Boone, Charles P. Gerba, M. Khalid Ijaz, Frank Y. Xu, Karl Krupp, Purnima Madhivanan, Kerry K. Cooper

## Abstract

Characterizing the household bacterial microbiome allows for a stronger understanding of the various microbes that a person is exposed to everyday in their home. Exploring household microbiomes in different countries around the world increases - our understanding of the impact cultural differences might have on niche microbial communities in the house. The goal of this study was to use shotgun metagenomics to characterize the microbiome for ten locations around the home in ten different houses from three different countries (Mysuru, India; Dubai, United Arab Emirates (UAE); and Tucson, United States of America (USA)). There was a significant difference in alpha diversity between the three countries (ANOVA, p<0.05) with homes in Mysuru, India showing significantly higher bacterial diversity compared to Dubai, UAE and Tucson, AZ, USA. Beta diversity analysis of the homes found that bacterial communities significantly differed between cities (PERMANOVA, p<0.01) and within cities by household locations (PERMANOVA, p<0.001). Locations such as underneath the toilet rim, bathroom and kitchen sinks had the highest levels of bacterial diversity across the three cities compared to other sampling areas. A core microbiome of Actinomycetes and Gammaproteobacteria was found in all homes in all three cities. Within each city, a core microbiome was identified at the species level within specific household locations in each city. Over 90% of bacterial taxa found in the homes were a part of the human-associated phyla Actinomycetes (eg. genera *Brevibacterium, Corynebacterium,* and *Microbacterium)*, Pseudomonadota (eg. genera *Acinetobacter, Moraxella, Pantoea, Paracoccus,* and *Psuedomonas)*, and Bacillota (genus *Streptococcus)*, which was comparable to previous studies. The household microbiome is variable in different locations in the house and on a global scale. Factors such as human activity, cultural practices, climate, and surface type and use may drive this diversity. Characterizing the household microbiome on a global scale allows for a better understanding of what drives microbial diversity, increasing our understanding of how microbial communities are shaped by the environment and how humans influence their dynamics, as well as any risks to human health that the built microbiome may potentially pose.

**Impact Statement:** This research contributes to the understanding of the built microbiome, specifically how it varies within the house, within cities, and across the globe. This can aid in our understanding of microbial dynamics in environments with heavy human influence and help develop and improve hygiene habits and products which are mindful of the existing microbiome.

**Data Summary:** DNA sequence data from this research is publicly available on the NCBI’s Sequence Read Archive under BioProject PRJNA1416920. Data was analyzed using python and R code. Analysis protocols and information on software versions, packages, and more can be found within the text and in the following github repository: https://github.com/carolinescranton01/Global_Household_Microbiome. The authors confirm all supporting data, code and protocols have been provided within the article or through supplementary data files.

## Introduction

The indoor microbiome is a complex and multifaceted environment influenced by a myriad of factors, both biotic and abiotic, that can have major impacts on human health^1–3^. The communities (bacteria, viruses, and eukaryotes) comprising the indoor microbiome are a joint result of direct and indirect factors including, but not limited to: the built environment location, the daily activities of the buildings’ occupants, and the surrounding external environment^3, 4^. Estimates show society has gradually spent increasing quantities of time indoors (from 65% upwards to 90% of human activity occurring in enclosed buildings in a typical 24-hour day), especially in geographic locales with higher degrees of socio-economic development and under the circumstances of the SARS-CoV-2 global pandemic^3–6^. Indoors, humans are key contributors to the surrounding microbiome where microorganisms can maintain viability for extended durations in aerosols or on fomites where active proliferation or persistence may occur for extended periods after introduction^3, 4, 7, 8^. Microbes inside may be transient or established, and survival varies between the materials and textiles comprising the household and is considerably influenced by the duration and frequency of sanitation and cleaning efforts of the occupants^4, 8, 9^. As a result, the transmission and preservation of microbes within the boundaries of the built environment they inhabit is complex and determined by a multitude of factors^3,4^.

Studies have shown that the persistence and presence of varying microbiological taxa in the indoor environment can provide both protective or adverse effects to occupants of the indoor space^4,7,9,10^. A crucial reservoir for household microbes is surface dust, comprising between 500-1,000 species according to some reports^9^. Analyses of the microflora of indoor dust have demonstrated the composition is complex, containing microbes and microbial secretions, particulates of human and animal origin (skin cells, dander, respiratory, hygiene), outdoor particulates (soil, plants), and indoor contributions (cooking, smoking, inorganics from cleaning products)^4, 8, 9, 11, 12^. Additionally, the introduction and persistence of indoor microbiota rely on physiological determinants including moisture, relative humidity, temperature, ultraviolet radiation, particle matter size, the pollutant and particulate indoor/outdoor ratio, and heat and ventilation^3, 4, 8, 13^. Studies have shown that buildings in more industrialized locations like major cities tend to have reduced soil-derived microbes in the indoor microbiome compared to those in rural areas^2, 8, 10^. Thus, the composition of the indoor microbiome is not universally consistent. Globally, the built environment varies in size, quality or components of building materials, ventilation, proximity to potential pollutants or greenspaces, and by geospatial climate^4, 6, 8^. Equivalently, occupant behavior disparities due to socio-economic factors and cultural dissimilarities influence the composition of the indoor microflora^4, 6, 9^. Therefore, understanding the indoor household microbiome and how it varies in different areas of the household, spatially between households located within the same region, and how it differs geographically between countries aid in our understanding of how the built microbiome develops and what risks it can potentially pose to household occupants.

In residential environments, previous indoor microbiome analyses of touch surfaces and interior dust deposition have demonstrated there may exist a relatively common core microbial phyla consisting of Pseudomonadota, Bacillota, and Actinomycetes^1,3, 7, 11, 13, 14^. The findings pose limitations as these phyla are heavily associated with the natural human microbiome of the skin, gut, oral cavity and secretions, and do not provide additional depth to other phyla which may shape the indoor bacterial microbiome irrespective to human occupants^15, 16^. Studies in the United States of America^7, 14^, India^13^, The United Kingdom, Greece^9^, and China^11^, as well as a global review^1^ examining the surfaces of kitchens, bathrooms and common living areas (living room or bedroom) frequently identified these “core” phyla alongside members of the Bacteroidota. The variability and diversity of microbial taxa within these phyla, however, were dependent upon the site or location of sampling where sites with moist environments typically harbored taxa different from those located on dry contact surfaces^3, 14^. Comparatively, locations frequently exposed to human associated sources harbored different taxa than those infrequently touched or those in proximity to outdoor air deposition^1, 7, 11, 13^. Not surprisingly, areas that lacked regular cleaning patterns typically showed greater microbial diversities^4^. Similarly, studies analyzing dust samples frequently identified the same “core” phyla across different geographic locations^1, 4, 9^. A study by Thompson et al. (2021) demonstrated differences in taxa based on geographic location where they observed a distinct country-specific microbiome variance between dust samples from United Kingdom and Greece^9^. A similar trend was observed in a study by Benton et al. (2023) that found floor dust in Nogales, Sonora, Mexico was more enriched with three different soil microbial genera when compared to dust from Tucson, Arizona in the United States despite a spatial separation of only 100 kilometers and similar climates^2^. However, differences in the study could also be attributed to socio-economic and structure of the built environment^2^.

It is important to understand the differences in microbial diversity across a range of household surfaces as well as geographic locations, and how factors such as socio-economic and cultural differences impact the microbiome. This provides information on the possible health implications for household occupants and presence of potential pathogens, which may be influenced by cleaning and hygiene habits as well as cultural differences. There are few studies assessing the household microbiome in our three study locations – some were conducted in the United States of America (USA), one was recently conducted in India, and none in the United Arab Emirates (UAE)^13, 14, 17^ This study assessed the household bacterial microbiome in three major cities (population >500,000) around the globe – Mysuru (Karnataka, India), Dubai (UAE), and Tucson (Arizona, USA). Ten different sampling locations were assessed in ten houses per city, focusing on the kitchen and bathroom. The microbiome was analyzed to determine dominant taxa across the locations and to identify a potential core microbiome, both within the cities and across the same household location in different cities across the world.

## Methods

### Household Sampling

Samples were collected from three geographic locations – Tucson (Arizona, USA – population est. 555,000^18^), Mysuru (Karnataka, India – population est. 3,000,000^19^) and Dubai (United Arab Emirates – population est. 3,800,000^20^). Participants (households) were recruited by local teams associated with the study, in January (Mysuru), February (Tucson), and September (Dubai). Households had between two and six adults and zero to three children residing there, with one to six bathrooms and two to six bedrooms. Households with pets were not excluded, but this was not considered in the presented analysis. Up to ten samples were collected from each household depending on availability, household construction, and participant willingness to provide access (**Supplementary Table 1**), for a total of 100 samples in Tucson and Mysuru and 89 samples in Dubai. Sampling locations included the kitchen counter surface, kitchen sink (surface and around but not inside the drain), coffee table surface, TV remote (all sides and buttons), showerhead, bathroom sink (surface and around but not inside the drain), and four locations on the toilet (all of which were standard western toilets) – the seat (outside and inside), rim of the toilet (top and outside), toilet bowl above the water, and the under the toilet rim (inside the bowl). Samples were collected with dual-end swabs (Becton Dickinson (Franklin Lakes, NJ, USA), BBL CultureSwab) that were dipped in a sterile detergent solution (0.15 M NaCl with 0.1% Tween 20)^21^ and then dragged across the sample surfaces for at least 15 seconds but no more than 30, rotating the swab during the sampling process. A specific surface area to swab was not specified for each household location, as there was too much variation between households and cities to standardize collection. However, the samples were taken from the entire surface of a particular targeted household sample location (i.e., corners, edges, center, etc.) to effectively sample all areas of the surface. Swabs were stored in their original sterile tubes and kept on ice in a cooler for less than 8 hours prior to storing at −20°C long term. Samples collected in Mysuru, India (n=100) were processed and sequenced on site at the Public Health Research Institute of India (PHRII) for the study. Samples from Tucson (n=100) were processed and sequenced in the Cooper lab (University of Arizona, Tucson, AZ), and samples collected in Dubai, UAE (n=89, as some households lacked the sampling locations of interest) were shipped on dry ice to the Cooper lab for processing and sequencing.

Prior to conducting the study, study design for Tucson and Dubai was reviewed by the University of Arizona’s Institutional Review Board (IRB), which determined that because no human identifying data was collected during the study it was exempt from an IRB. The study design for households in India was approved by the Public Health Research Institute of India’s IRB Committee prior to sampling.

### DNA Extraction

DNA extraction from each sample included removing one of the two swabs and conducting extraction using the DNeasy Powersoil Pro Kit (Qiagen) per the manufacturer’s instruction with one modification – samples were heated at 65°C for 10 minutes after adding the first lysis buffer (CD1)^22^. DNA concentrations were measured on the Qubit fluorometer using the dsDNA High-Sensitivity Assay Kit (Invitrogen (Carlsbad, CA, USA)). If the DNA concentration was deemed too low by the Qubit assay (less than 0.0005 ug/mL), the second swab was used for DNA extraction to increase yield.

### Oxford Nanopore Sequencing

Sequencing libraries in the Cooper laboratory were prepared using the Oxford Nanopore Rapid PCR Barcoding Kit 24 V14 (Oxford Nanopore Technology, SQK-RPB114.24, protocol version RBK_9176_v114_revK_27Nov2022) per the manufacturer’s instructions. Sequencing libraries prepared at PHRII used the Oxford Nanopore Rapid Barcoding Kit 96 V14 (Oxford Nanopore Technology, SQK-RBK114.96, protocol version RBK_9176_v114_revK_27Nov2022) according to the manufacturer’s protocols. This PCR-free library preparation kit was used as there were no thermocyclers available on-site at PHRII, so samples were sequenced multiple times to increase sequencing yeild. All samples were sequenced using Oxford Nanopore Technology (ONT) on MinION Mk1B devices (Oxford Nanopore Technology) with all libraries being sequenced using R10.4.1 flow cells (Oxford Nanopore Technology). Libraries were sequenced multiple times in both locations to increase the data yield from each sample. Negative control samples were not collected or sequenced.

### Taxonomic Analysis

Reads were generated using multiple sequencing runs to increase data yield and concatenated across runs to form fastq files for each sample in the ten sample locations within the houses, in the three cities (ten samples per house, ten houses per city; n=100). Additionally, the reads from the same location across the ten houses in each city (i.e. all bathroom sinks in Tucson) were concatenated to create composite samples for each household location in each city in order to assess the variability of each location overall. Reads were then analyzed for taxonomic composition using Kraken2^23^ software (v2.3.1), using a database that contained all reference genomes from the NCBI database as well as all non-redundant sequences (nr database, as of February 17, 2023), which was created by the Cooper laboratory following the kraken2-build protocol for Kraken2^23^. Kraken2 was run on all concatenated fastq files (non-composites and composites) with flags to specify the database location, and what to name the output and report. BIOM-format files were generated with the kraken_biom software (v1.2.0)^24^ and analyzed as phyloseq objects using the R package phyloseq (v1.52.0)^25^. After importing the data into R (v4.3.5) using RStudio (v2024.04.0+735), it was filtered to only include bacterial taxa (which accounted for approximately 70% of the reads) – this was done by removing all taxonomic assignments outside of the domain Bacteria. The remaining 30% of reads primarily belonged to the phyla Chordata and Arthropoda, which were excluded from the analysis. This filtered phyloseq object was used for the bacterial taxonomic, alpha diversity, beta diversity, and core microbiome analyses, as described below using the following R packages: phyloseq, microbiome (v1.30.0)^26^, microViz (v0.12.7)^27^, and vegan (v2.7.3)^28^.

### Bacterial Diversity Analysis

To assess the alpha diversity of both the geographical and household sampling location, different alpha diversity parameters were calculated (Shannon, Simpson, Chao, Fisher, and observed species) using the microbiome package in R. Analysis of variance (ANOVA) tests were used to determine significant differences in the alpha diversity of the samples within the three cities overall and between the ten household locations within each city. The Bray-Curtis dissimilarity index was used to determine the beta diversity of the different samples in the study. Permutational multivariate analysis of variance (PERMANOVA) tests were conducted on the Bray-Curtis distances to determine statistical significance between the household locations, geographic locations, and the combination of both household and geographic location.

Relative abundance of the top taxa was calculated at the phylum, class, order, family, and genus levels using the aggregate_taxa command from the microbiome package. Composite data for each household location within each city was used in this analysis when looking at the ten household locations individually. The entire dataset excluding composite samples was used to determine the top taxa across the three cities as a whole. The core microbiome was analyzed using the aggregate_taxa and aggregate_rare functions in the microbiome R package for each city individually, across each household location within a particular city, and across all three cities overall, at a prevalence level of 90% (taxa must be present in 90% of samples) and a detection level of 1% (taxa must be detected at a relative abundance of 1% or more in the sample to be included) for higher taxonomic orders (phyla, class order, and family) and 0.1% for the species level. Three species detected in the core microbiome analysis were known to share a large amount of their genomes – *Salmonella enterica, Escherichia coli*, and *Shigella flexneri.* Reads mapped to these samples using kraken2 were identified through NCBI taxIDs in the kraken reports and were mapped to reference genomes for each species to ensure that these species were identified correctly.

### Data Visualization and Availability

RStudio and Microsoft Excel were used to subset and reorganize the data. Results were represented using visual plots, generated using ggplot2 (v4.0.2)^29^ in R using RStudio. The packages RColorBrewer (v1.1.3)^30^, ggpubr (v0.6.3)^31^, ggplot2 were used to visualize the relative abundance of taxa, plot alpha diversity metrics as box-and-whisker plots, and to plot a principle components analysis of the taxonomic makeup of the samples using Bray-Curtis distances. The packages tidyr (v1.3.2)^32^, data.table (v1.18.2.1)^33^, dplyr (v1.2.0)^34^, openxlsx (v4.2.8.1)^35^, and writexl (v1.5.4)^36^ were used to restructure data (add or change metadata, filter subsets of data, transpose data) throughout the analysis. Code used for data analysis can be found on GitHub (https://github.com/carolinescranton01/Global_Household_Microbiome). All raw sequence data has been submitted to NCBI’s Sequence Read Archive and is available under the BioProject accession PRJNA1416920.

## Results

### Global

The goal of this study was to examine the diversity of the household bacterial microbiome between different countries around the world, and therefore households from three cities with at least 500,000 residents were sampled including Tucson, AZ, USA (∼575,000 people); Dubai, UAE (∼3.65 million people); and Mysuru, India (∼3 million people). Looking at the household in general, alpha diversity of the overall household for each city found there was a significant difference based on the location, particularly with households in Mysuru having higher diversity compared to Tucson or Dubai regardless of the diversity index used (Figure 1A; **Supplementary Figure 1A-C**). Beta diversity based on the Bray-Curtis dissimilarity index among all samples found that samples from Dubai and Tucson clustered the tightest together with overlap between the two cities, whereas household samples from Mysuru were spread out, indicating more variation between the homes in this city (Figure 1B). PERMANOVA analysis revealed significant differences by geographic location (p<0.001), household location individually (p<0.001), and when household and geographic location were considered together (p<0.001).

**Figure 1.**
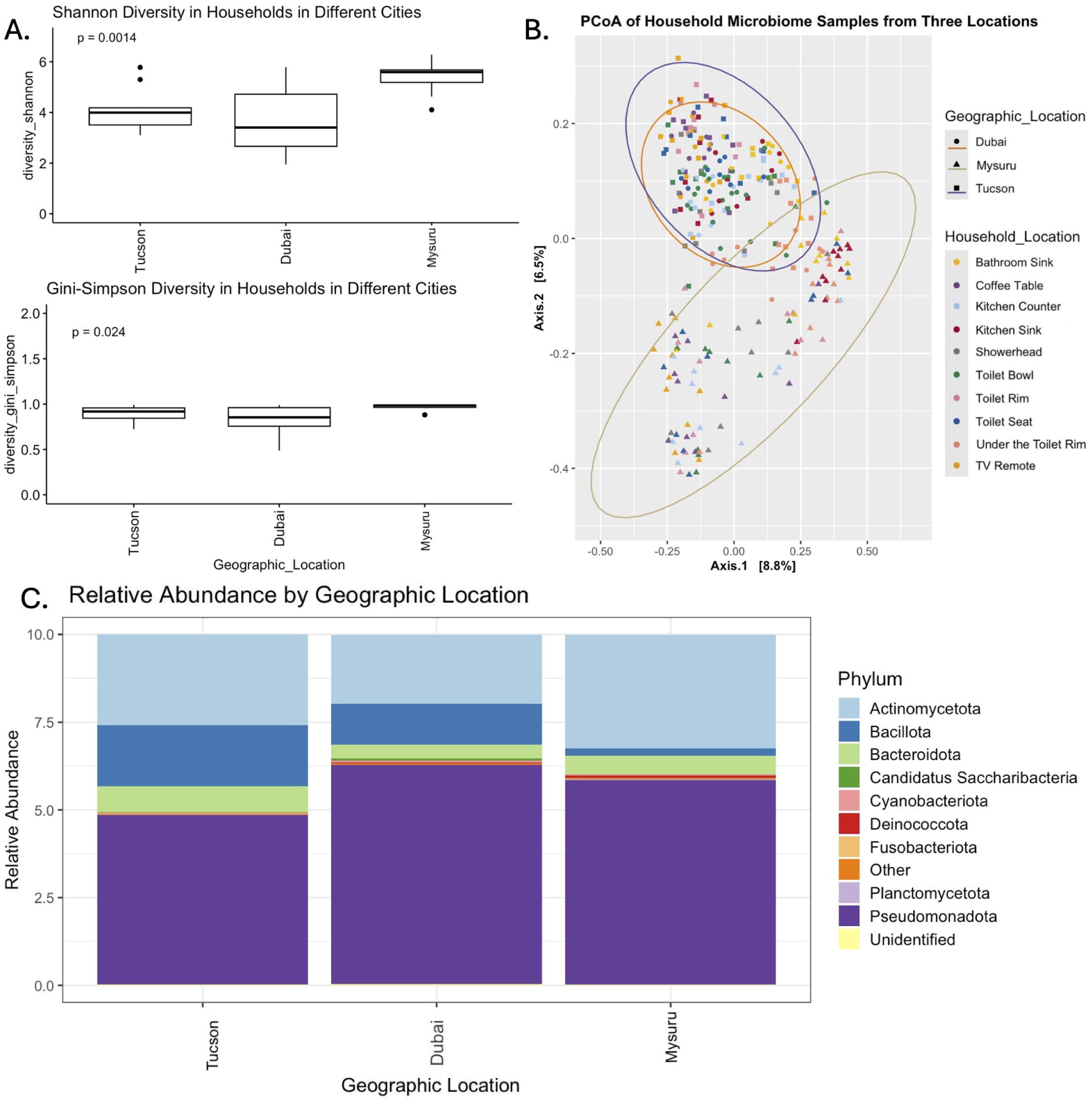
Taxonomic composition of the household microbiome overall between the three study locations. A: Shannon and Simpson diversity indices (alpha diversity) for the three study locations overall between the cities – Shannon diversity was highly significant (ANOVA, p<0.01) and Gini-Simpson diversity was significant (ANOVA, p<0.05). B: Principal coordinates analysis of Bray-Curtis distances (beta diversity) for all samples in the study. Samples are colored by location within the house, point shapes correspond to the geographic location, and ellipses are drawn around clusters by geographic location. Significant differences were found between both geographic location, household location, and the interaction term of the two (PERMANOVA, p>0.001). C: The relative abundance of the top 10 most abundant phyla per geographic location.

The phylum Pseudomonadota had the highest abundance in the household for each of the three cities, making up 48% to 63% of the bacterial microbiome, with Dubai having the highest levels and Tucson having the lowest levels. The second highest phylum among the three cities was Actinomycetota, with Mysuru having the highest levels (29% of the bacterial microbiome) and Dubai the lowest (21%), while Bacillota was highest in Tucson (7%) and lowest in Dubai (5%) (Figure 1C). At the class level, Gammaproteobacteria was the most abundant in all three cities, with Dubai having the highest levels and Mysuru the lowest (**Supplementary Figure 2A**). The order Moraxellales was the most abundant in Dubai and was also the dominant order in Tucson, but in Mysuru households Micrococcales was the most abundant identified order (**Supplementary Figure 2B**). Tucson households had families Moraxellaceae, Mycobacteriaceae, and Enterobacteriaceae as the three most abundant bacterial families, whereas Moraxellaceae and Erwiniaceae were most abundant in Dubai households. Mysuru households had less Moraxellaceae compared to the other two cities, but it was still the most abundant followed by Enterobacteriaceae and Paracoccaceae (**Supplementary Figure 2C**). The household core microbiome for all three cities included the Pseudomonadota phyla and at the class level Acintomycetes and Gammaproteobacteria. No core microbiome was found at any lower taxonomic levels in Dubai or Tucson, or across all three cities, but in Mysuru households there were five species found in all samples at a 0.1% abundance or greater– *Escherichia coli, Francisella halioticida, Moraxella osloensis, Salmonella enterica, and Shigella flexneri* (Table 2).

Next, we explored the actual sampling locations within the household for each of the three cities to gain a better understanding of the specific household microbiome composition. The Shannon diversity index found there were significant differences between the household sampling sites in households in all three cities. Tucson households had the highest level of diversity under the toilet rim, in the kitchen sink, and in the toilet bowl, whereas the TV remote, under the toilet rim, and inside the toilet bowl had the highest bacterial diversity in Dubai households. In Mysuru households, under the toilet rim, in the kitchen sink and the kitchen counter had the highest levels of diversity. Overall, under the toilet rim has one of the highest levels of bacterial diversity (ranging from 3 to 6.5 on the Shannon index) among the household samples regardless of the country. The lowest level of bacterial diversity varied by country with the toilet seat and kitchen sink in Dubai, TV remote and showerhead in Mysuru, and kitchen counter and showerhead in Tucson all having lower levels of diversity compared to the other household locations in those cities (Figure 2A; **Supplementary Figure 3A-D**). The Bray-Curtis PCoA dissimilarity distances of the composite samples for each household location for the three cities found that none of the cities clustered very tightly, but Dubai and Tucson samples clustered closer together compared to samples from Mysuru. PERMANOVA analysis showed significant differences in the taxonomic profile of the composite samples by geographic location (p<0.001), but not by household location (p=0.533). Despite this, some clustering based on household location can be observed between the three cities such as under the toilet rim, TV remotes, and bathroom sinks, all of which clustered together regardless of geographic location (Figure 2B).

**Figure 2.**
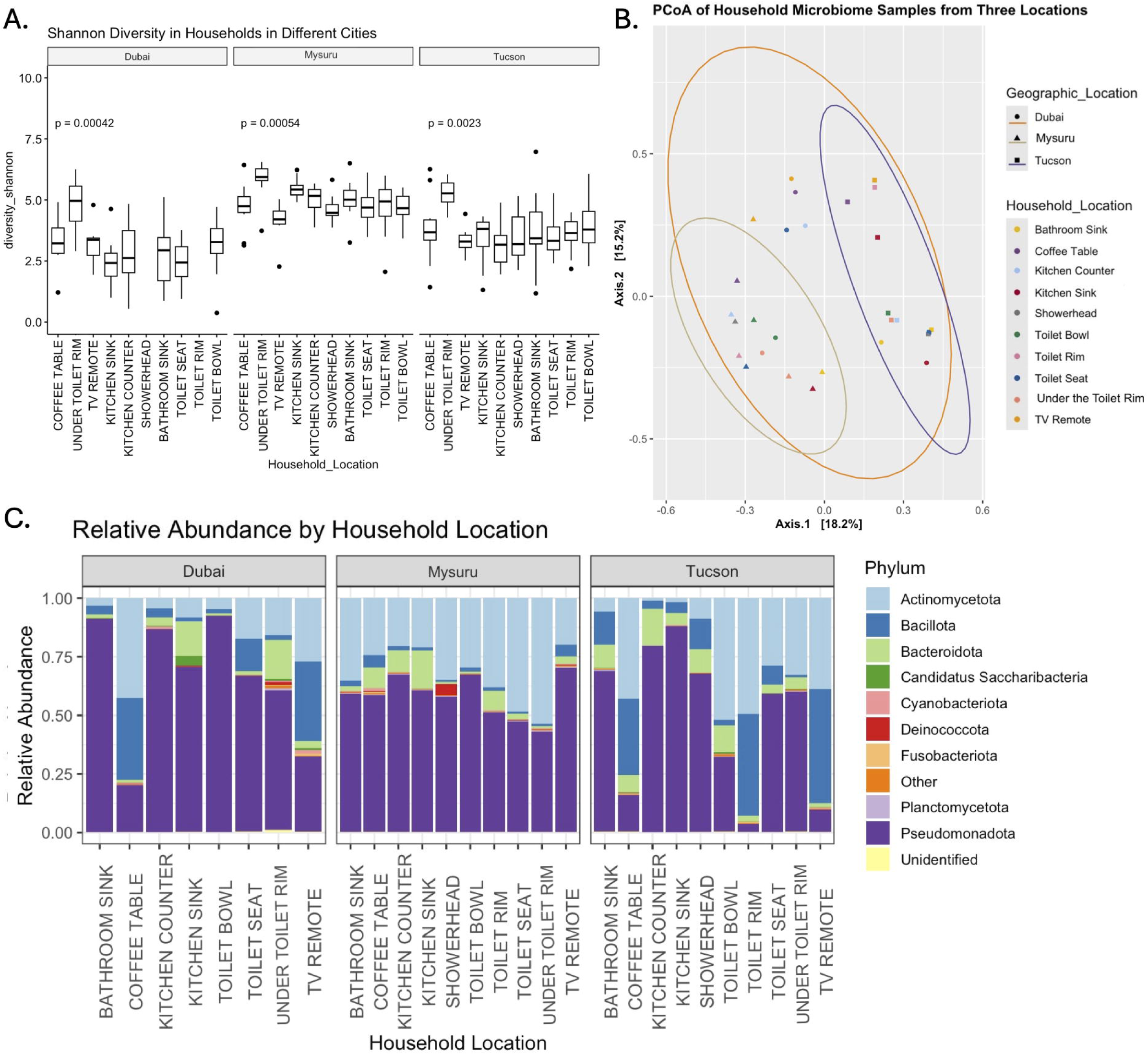
Taxonomic composition of the ten different household locations within each house, from the three study locations. A: Shannon diversity index (alpha diversity) for the three study locations overall, which only significantly differed in Mysuru (ANOVA, p<0.01). B: Principal coordinates analysis of Bray-Curtis distances (beta diversity) for composite samples (all household locations per geographic location merged, ie. All kitchen sinks in Mysuru) in the study. Samples are colored by location within the house, point shapes correspond to the geographic location, and ellipses are drawn around clusters by geographic location. Significant differences were found between geographic locations (PERMANOVA, p<0.01) but not household locations (PERMANOVA, p=0.533). C: The relative abundance of the top 10 most abundant phyla per household location in each city.

The taxonomic composition of the different household samples from Mysuru all contained the phylum Pseudomonadota at the highest abundance with Actinomycetota being the second most abundant phylum. The coffee table, kitchen counter, kitchen sink, and toilet rim samples had higher abundance of Bacteriodota in Mysuru and Tucson compared Dubai. In Dubai the bathroom sink, kitchen counter, kitchen sink, toilet bowl, toilet seat, and under the toilet rim all had high abundance of the phylum Pseudomonadota, whereas the coffee table and TV remote had higher abundances of Actinomycetota and Bacillota. Household samples from Tucson were similar to Dubai, as the bathroom sink, kitchen counter, kitchen sink, showerhead, toilet seat, and under the toilet rim had the phylum Pseudomonadota as the highest abundance, but in Tucson, the toilet bowl samples were over 50% Actinomycetota. Furthermore, the coffee table, toilet rim, and TV remote in Tucson samples were dominated by the high abundance of the phyla Actinomycetota and Bacillota (Figure 2C).

At the class level, we found a consistent distribution in Mysuru of the most abundant bacterial classes, which were Gammaproteobacteria, Actinomycetes, and Alphaproteobacteria. However, the TV remote and toilet bowl had much higher levels of Gammaproteobacteria, and under the toilet rim, the toilet seat, and the toilet rim had higher levels of Actinomycetes. In contrast, both Dubai and Tucson had more variable composition with higher levels of Gammaproteobacteria in numerous samples and higher abundance of Bacilli on the coffee table, TV remote, and in Tucson, on the toilet rim. Overall, at the class level, taxonomic profiles for the various household samples from Dubai were much closer to Tucson than Mysuru. In all cities, most of the bacteria (85% to 99%, depending on the sample) were part of the top ten classes in the samples, as denoted by low percentage of grouped ‘other’ category (**Supplementary Figure 4A**).

The taxonomic profile at the order level started to demonstrate a separation between Tucson and Dubai samples, as the bathroom sink, kitchen sink, toilet bowl, and toilet seat in Dubai were dominated by higher Moraxellales. Although Tucson also had high levels on the kitchen counter, bathroom sink, toilet seat, and showerhead, it also had higher abundance of other bacterial orders. The kitchen sink in Tucson had a high level of diversity, including Enterobacterales, Moraxellales, and Pseudomonadales, compared to Dubai that was dominated by Moraxellales and Flavobacteriales. The toilet bowl varied greatly between the three cities, where in Tucson it was dominated by Mycobacteriales, but in Dubai and Mysuru Moraxellales was most abundant, however Mysuru had a greater variety of additional taxa in that location as shown by the larger ‘other’ category which was consistently found in each household location. Again, Mysuru household samples taxonomic profiles were quite different from the other two cities with a more consistent distribution of the different orders. Each city had one location in the house that were dominated by Enterobacterales, but it varied in each city including the kitchen counter in Dubai, TV Remote in Mysuru, and kitchen sink in Tucson (**Supplementary Figure 4B**). However, at the family level there was only elevated *Enterobacteriaceae* in Tucson households in the kitchen sink and Mysuru households on the TV remote. *Moraxellaceae* was the dominant bacterial family in numerous household locations in the three cities including the bathroom sink, kitchen sink, toilet bowl, under the toilet rim, TV remote, and toilet bowl in Dubai, the toilet bowl and kitchen sink in Mysuru, and bathroom sink, kitchen counter, shower head and toilet seat in Tucson households (**Supplementary Figure 4C**).

Examining the core microbiome at the species level for each of the different household locations of the three countries supported the previous analysis that Mysuru had the largest and most diverse core microbiome among the three cities at each household location. Dubai and Tucson had slightly smaller core microbiomes at the different locations across the household, composed of zero to four species, while Mysuru’s core microbiome across each location ranged from four to nine species (Table 2). The overall core microbiome at the family level across the three cities found a core microbiome in Mysuru only, composed of *Enterobacteriaceae* and *Paracoccaceae* (Table 1).

**Table 1.** Core Microbiota in each location.

| **Taxonomic Level** | **Dubai** | **Mysuru** | **Tucson** | **All Locations** |
| --- | --- | --- | --- | --- |
| Phylum<sup>1</sup> | Pseudomonadota | Actinomycetota<br>Pseudomonadota | Actinomycetota<br>Bacillota<br>Pseudomonadota | Pseudomonadota |
| Class<sup>2</sup> | Gammaproteobacteria | Actinomycetes<br>Alphaproteobacteria<br>Betaproteobacteria<br>Gammaproteobacteria | Actinomycetes<br>Gammaproteobacteria | Actinomycetes<br>Gammaproteobacteria |
| Order<sup>3</sup> | None | Burkholderiales<br>Enterobacterales<br>Hyphomicrobiales<br>Micrococcales<br>Mycobacteriales<br>Rhodobacterales | None | None |
| Family<sup>4</sup> | None | Enterobacteriaceae<br>Paracoccaceae | None | None |

<sup>1</sup>Phylum level, prevalence = 90%, detection = 1% <sup>2</sup>Class level, prevalence = 90%, detection = 1% <sup>3</sup>Order level, prevalence = 90%, detection = 1% <sup>4</sup>Family level, prevalence = 90%, detection = 1%

**Table 2.**
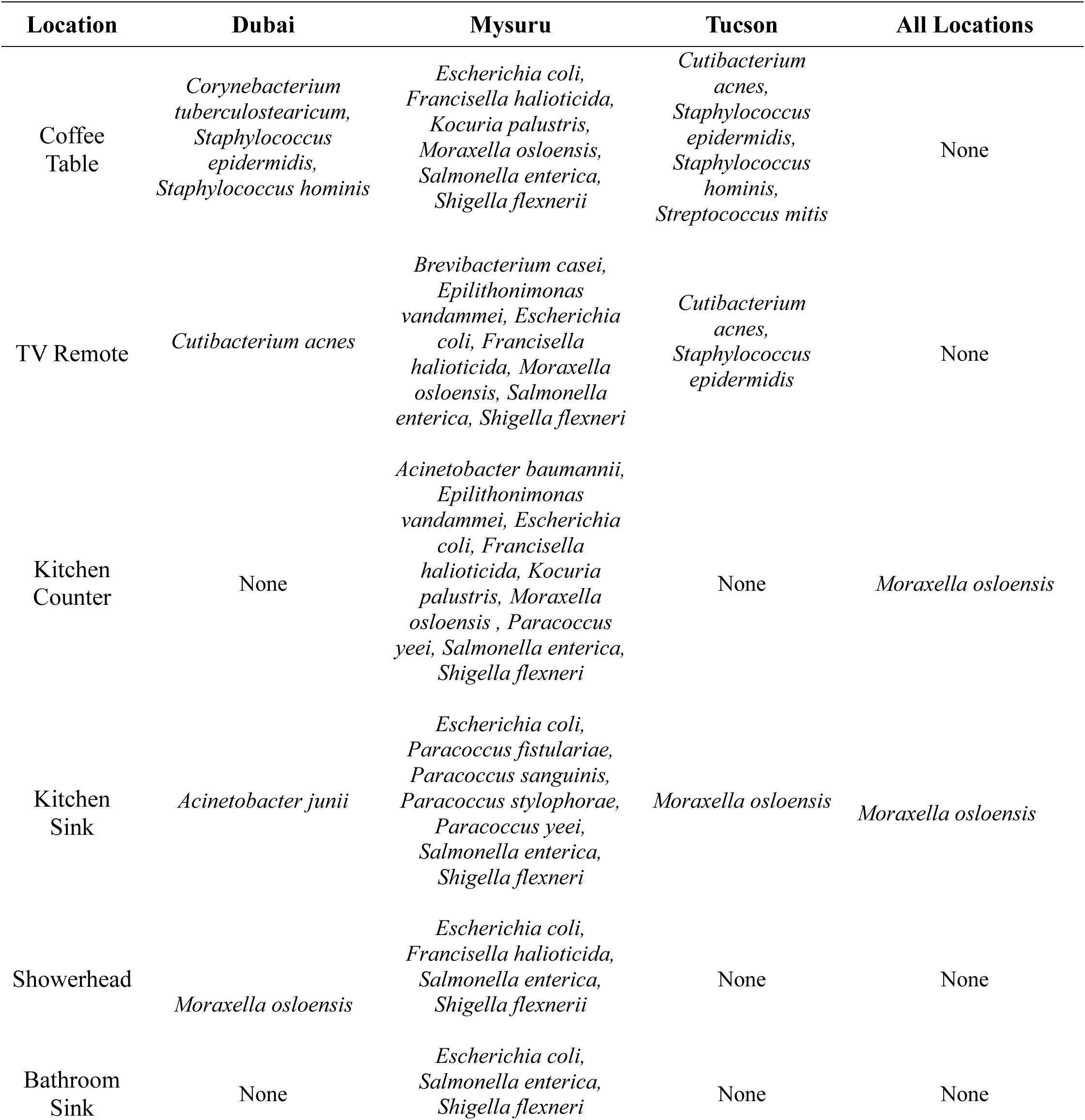
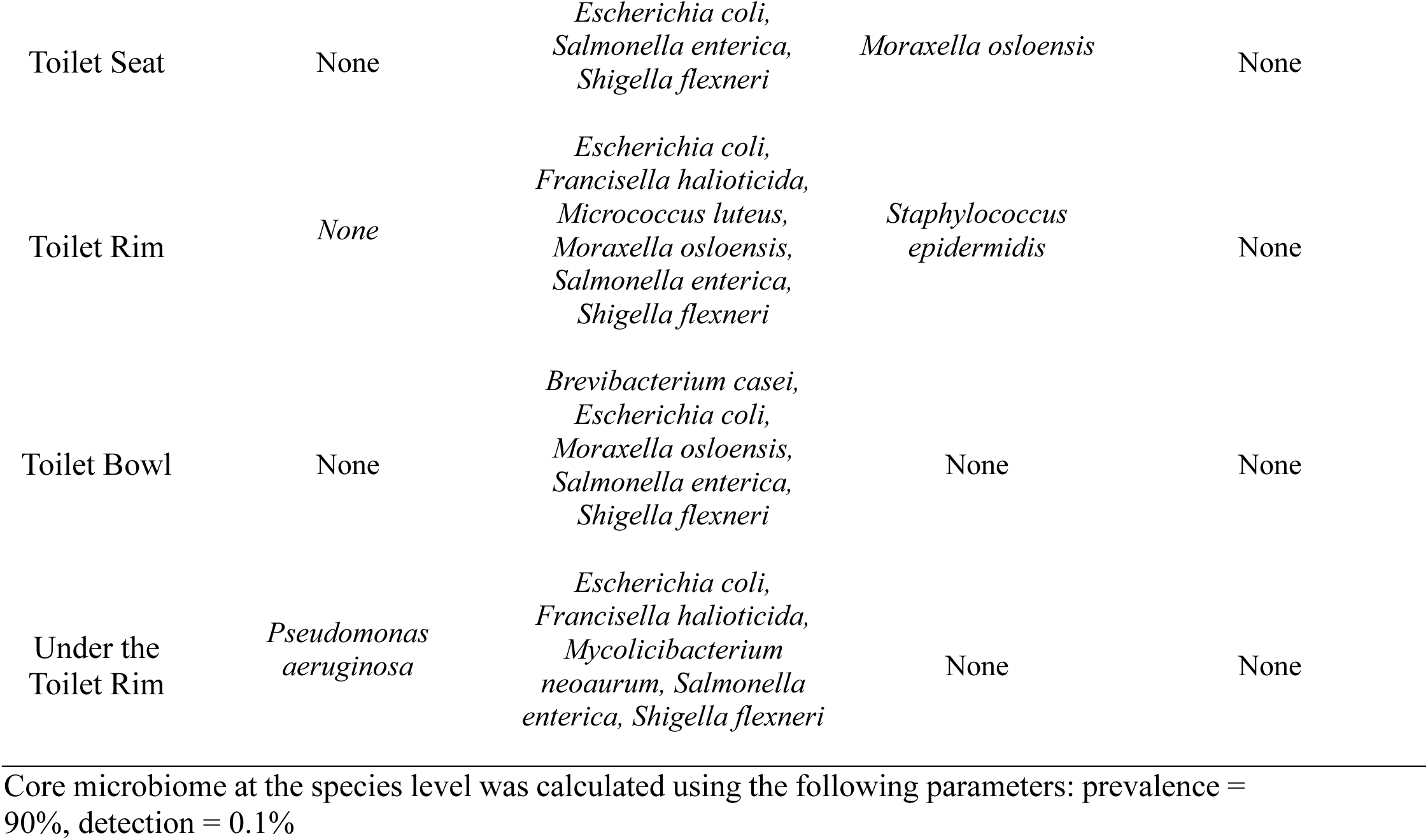
Core microbiome at the bacterial species level for each household sampling site in each location.

| Location | Dubai | Mysuru | Tucson | All Locations |
| --- | --- | --- | --- | --- |
| Coffee Table | *Corynebacterium tuberculostearicum*,<br>*Staphylococcus epidermidis*,<br>*Staphylococcus hominis* | *Escherichia coli*,<br>*Francisella haliotica*,<br>*Kocuria palustris*,<br>*Moraxella osloensis*,<br>*Salmonella enterica*,<br>*Shigella flexnerii* | *Cutibacterium acnes*,<br>*Staphylococcus epidermidis*,<br>*Staphylococcus hominis*,<br>*Streptococcus mitis* | None |
| TV Remote | *Cutibacterium acnes* | *Brevibacterium casei*,<br>*Epilithonimonas vandammei*, *Escherichia coli*, *Francisella haliotica*, *Moraxella osloensis*, *Salmonella enterica*, *Shigella flexneri* | *Cutibacterium acnes*,<br>*Staphylococcus epidermidis* | None |
| Kitchen Counter | None | *Acinetobacter baumannii*,<br>*Epilithonimonas vandammei*, *Escherichia coli*, *Francisella haliotica*, *Kocuria palustris*, *Moraxella osloensis*, *Paracoccus yeei*, *Salmonella enterica*, *Shigella flexneri* | None | *Moraxella osloensis* |
| Kitchen Sink | *Acinetobacter junii* | *Escherichia coli*,<br>*Paracoccus fistulariae*,<br>*Paracoccus sanguinis*,<br>*Paracoccus stylophorae*,<br>*Paracoccus yeei*,<br>*Salmonella enterica*,<br>*Shigella flexneri* | *Moraxella osloensis* | *Moraxella osloensis* |
| Showerhead | *Moraxella osloensis* | *Escherichia coli*,<br>*Francisella haliotica*,<br>*Salmonella enterica*,<br>*Shigella flexnerii* | None | None |
| Bathroom Sink | None | *Escherichia coli*,<br>*Salmonella enterica*,<br>*Shigella flexneri* | None | None |
| Toilet Seat | None | *Escherichia coli*,<br>*Salmonella enterica*,<br>*Shigella flexneri* | *Moraxella osloensis* | None |
| Toilet Rim | None | *Escherichia coli*,<br>*Francisella haliotica*,<br>*Micrococcus luteus*,<br>*Moraxella osloensis*,<br>*Salmonella enterica*,<br>*Shigella flexneri* | *Staphylococcus epidermidis* | None |
| Toilet Bowl | None | *Brevibacterium casei*,<br>*Escherichia coli*,<br>*Moraxella osloensis*,<br>*Salmonella enterica*,<br>*Shigella flexneri* | None | None |
| Under the Toilet Rim | *Pseudomonas aeruginosa* | *Escherichia coli*,<br>*Francisella haliotica*,<br>*Mycobacterium neoaurum*,<br>*Salmonella enterica*,<br>*Shigella flexneri* | None | None |

Core microbiome at the species level was calculated using the following parameters: prevalence = 90%, detection = 0.1%

### Mysuru, India

Next, we examined the household microbiomes within each specific geographic location to gain a better city-specific perspective of the household microbiome. The beta diversity of the various households in Mysuru found that there was a large amount of taxonomic variation between the household locations with slight clustering of samples based on household sampling location, such as TV remote and coffee table, and toilet seat and toilet rim. The toilet bowls, bathroom sinks, kitchen counters, and showerheads were more dispersed among the various household samples (Figure 3A). The beta diversity of samples significantly differed between each household location (PERMANOVA, p<0.001). All locations except the toilet bowls have at least 40% of the relative abundance as “other” bacterial genera (genera which were not part of the top ten most abundant across all Mysuru samples). All the household sampling locations except under the toilet rims and the showerheads contained at least 5% relative abundance of *Acinetobacter* with the toilet bowl having close to 50%. Additionally, levels of *Paracoccus*, *Epilithonimonas, Moraxella,* and *Mycolicibacterium* were slightly elevated in several household locations (Figure 3B).

**Figure 3.**
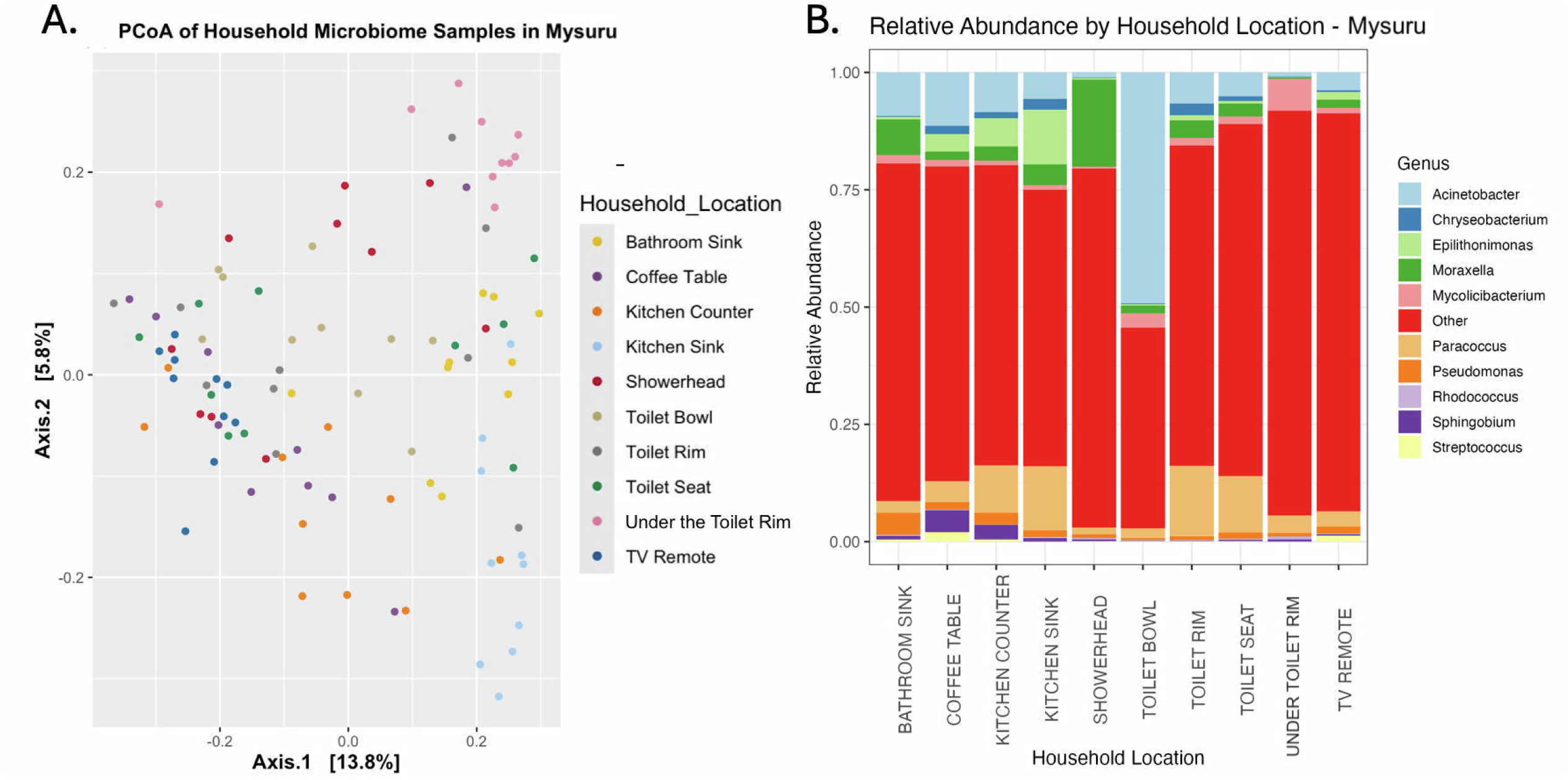
Taxonomic composition of samples from Mysuru, India. A: Principal Coordinates analysis (PCoA) of Bray-Curtis distances for all samples collected in Mysuru. Points are colored by household location. Significant differences in taxonomic composition were found between household locations (PERMANOVA, p<0.001). B: Relative abundance of the top 10 taxonomic genera from each household location.

### Dubai, UAE

PERMANOVA analysis found significant differences in the beta diversity of the different household locations in Dubai (p<0.001). The toilet bowl, toilet seat, and under the toilet rim samples clustered more with each other compared to other samples, as did the coffee table and TV remote samples (Figure 4A). Comparing the relative abundance of genera in the different household locations found *Moraxella* was the dominate bacterial genus on the bathroom sink, kitchen sink, toilet bowl, and toilet seat, while *Acinetobacter* was present in all the locations in an abundance of 2% or higher, but was dominant in the bathroom sink, kitchen sink, toilet bowl and toilet seat (Figure 4B). *Streptococcus* was found at slightly higher levels on the TV remote than in other locations across Dubai households.

**Figure 4.**
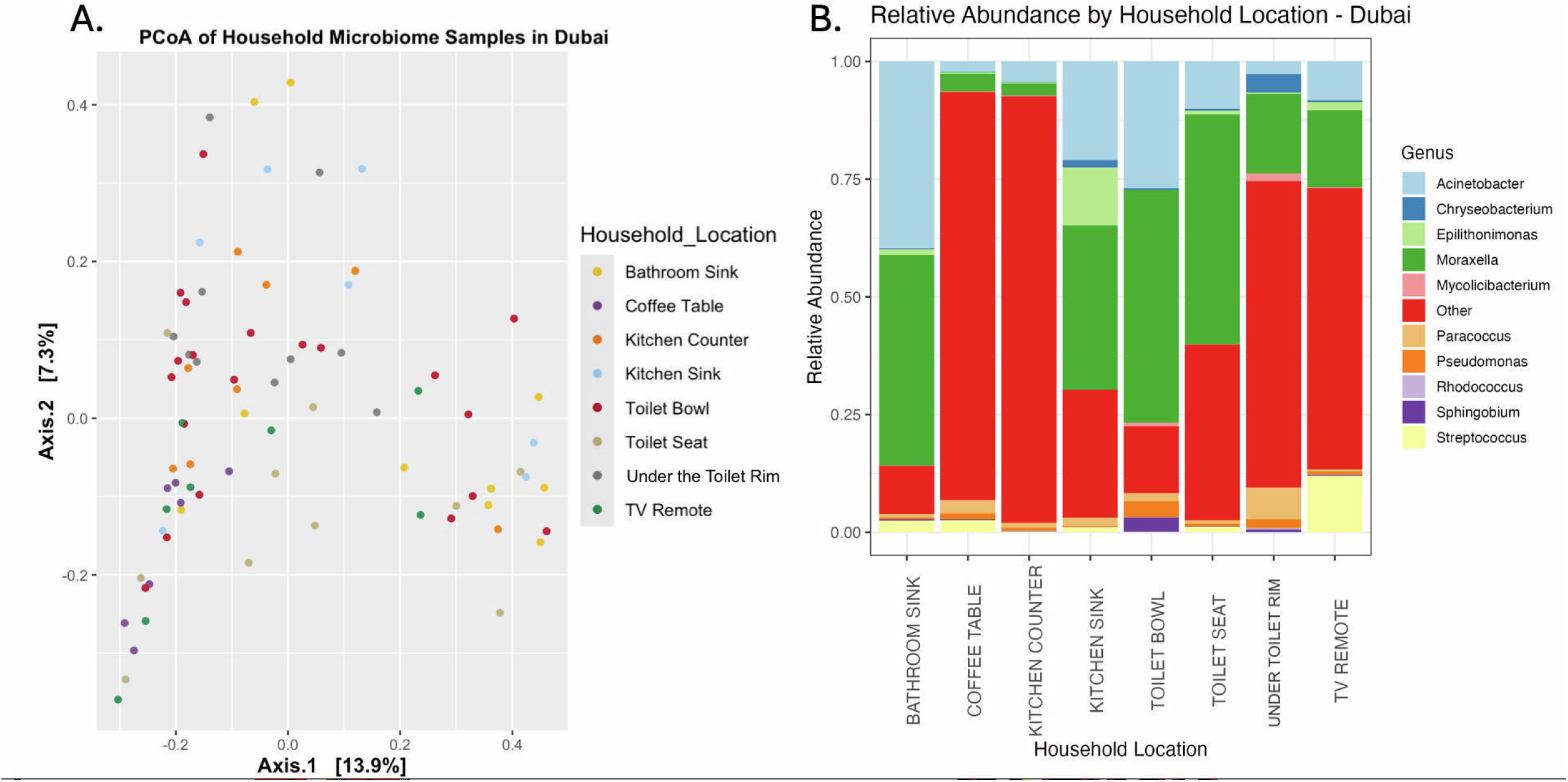
Taxonomic composition of samples from Dubai, UAE. A: Principal Coordinates analysis (PCoA) of Bray-Curtis distances for all samples collected in Dubai. Points are colored by household location. Significant differences in taxonomic composition were found between household locations (PERMANOVA, p<0.001). B: Relative abundance of the top 10 taxonomic genera from each household location.

### Tucson, Arizona, USA

Compared to Dubai and Mysuru, the beta diversity of the microbiome of specific household locations clustered more tightly and were highly significant based on the sampling areas in Tucson (PERMANOVA, p<0.001). The kitchen counters, kitchen sinks and bathroom sinks were clustered together, as were the TV remote, toilet rim, toilet seat, and coffee table (Figure 5A). The taxonomic makeup varied across locations in Tucson – in the bathroom sink, on the showerhead, and toilet seat, *Moraxella* was the most abundant genus, accounting for roughly 50% of the taxa on these locations. *Acinetobacter* was found to make up over 50% of the composition of the kitchen counter samples. Similarly to Dubai, *Streptococcus* was found at low levels across all samples but was notably more abundant on the TV remote. *Mycolicibacterium* was found at higher levels in the toilet bowl and under the toilet rim, similarly to Mysuru. Samples from Tucson appeared to show a larger amount of diversity in the most abundant genera across the different household locations compared to Mysuru or Dubai (Figure 5B).

**Figure 5.**
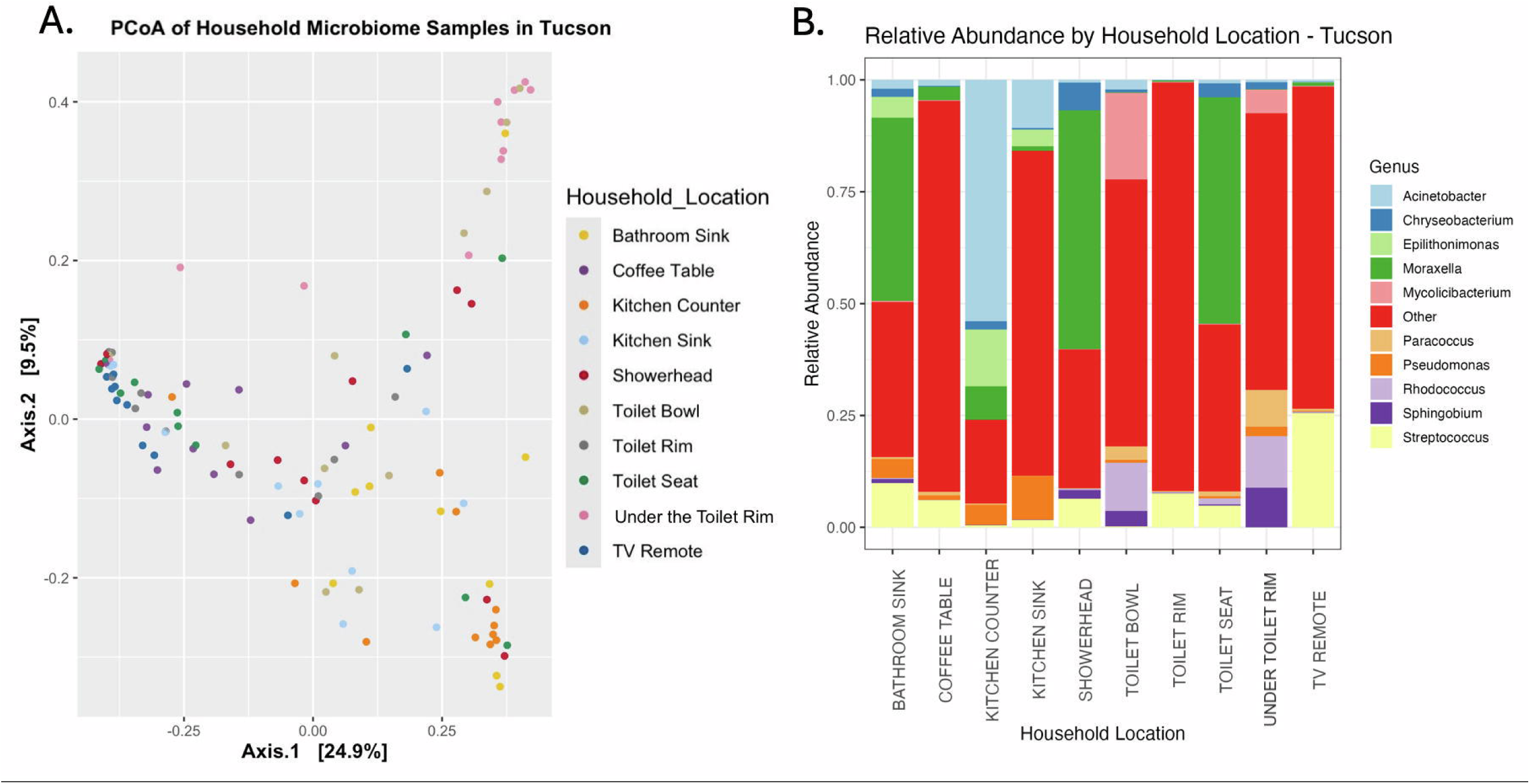
Taxonomic composition of samples from Tucson, Arizona, USA. A: Principal Coordinates analysis (PCoA) of Bray-Curtis distances for all samples collected in Mysuru. Points are colored by household location. Significant differences in taxonomic composition were found between household locations (PERMANOVA, p<0.001). B: Relative abundance of the top 10 taxonomic genera from each household location.

## Discussion

The composition of the bacterial microbiome varied significantly in all household sampling locations, between the houses, and across the three cities. The most common taxa found in this study were in the phyla Pseudomonadota (Proteobacteria), Actinomycetota (Actinobacteria), and in Tucson and Dubai (lesser so in Mysuru), Bacillota (Firmicutes), similar to previous global studies in built environments, whether they were houses, businesses, or schools^12, 37, 38^. These taxa are highly associated with the human microbiome – Actinomycetota dominates the respiratory, skin, and gut microbiome, while Bacillota is found abundantly in the mouth and respiratory tract, along with Psuedomonadota^14–16, 39, 40^. Many spore-forming and soil-associated bacteria are in the Bacillota and Psuedomonadota phyla and have been associated with dust in desert areas and within households^39, 41, 42^. Previous research has found these phyla to be common in indoor environments, along with Bacteriodota^1, 3, 7, 11, 13, 14^, which was found at about a 5% relative abundance in each of the geographic locations in this study. Considering that the microbiome of built environments has been shown to shift from environmental microbes to human-related microbes as the length of inhabitance increases^43^, the high prevalence of human-associated bacteria in the microbiome in this study and others is consistent with this information. We also identified several pathogenic species of bacteria, such as *Escherichia coli* and *Actinobacter baumanii* in the core microbiome of different household locations across the cities. It is important to note that the presence of pathogen DNA on household surfaces does not signify viable infectious organisms, and more work should be done to assess the risk to human health when pathogenic bacterial DNA is identified on household surfaces. Knowing that globally, the microbiome of the built environment is dominated by human-associated taxa, studying their roles and genes may aid our understanding of how the microbiome of the built environment impacts human health as these microbes can live on and within human.

Although many of the same phyla are found across the globe, our study found a high level of diversity in both the types and proportions of different taxa particularly at the lower taxonomic levels. Often, the same few classes, orders, families, or genera dominated the microbiome, but at different proportions in the three cities and/or household locations. At the class level, human-associated Actinomycetes and Gammaproteobacteria were found to be core members across all samples from the study, which is similar to what was found in other studies from the United States, India, and China^1, 7, 11^. Awasthi et al. conducted the only other household microbiome study in India to date that examined homes in northern India compared to our study in Southern India and still found similar results to ours. Both studies found at the phyla level, Psuedomonadota and Actinomycetota were the most abundant, while at the species level, *Escherichia coli* and *Moraxella osloensis* were found throughout the household as core microbiome members^13^. These two patterns were not observed in the Tucson and Dubai data. Another study from the United States found similar bacterial families to those in our Tucson data being most abundant in the kitchen, such as *Moraxellaceae* (highly abundant on Tucson kitchen counters but not the other cities in this study) and *Streptococcaceae*, commonly associated with the human skin microbiome, on the TV remote (found more often in Tucson than Dubai or Mysuru)^13, 14^. In contrast, another study that examined the household microbiome in five countries in Europe found different core and abundant taxa, including families like *Yersiniaceae* and genera such as *Bacillus* and *Enhydrobacter*, whereas these taxa were hardly if at all detected in our analysis^38^. One potential driver behind this difference is that the researchers in Europe used 16S rRNA amplicon sequencing while we used a shotgun metagenomics approach – it is known that there is some variation in sequencing results between these two methods, or potentially the age of the city and household or population density differences which may impact the built microbiome^44^. There are clear, significant differences in the taxonomic makeup of the microbiome across geographic space, and there are many potential drivers of these differences. When clustering the samples based on Bray-Curtis distance, samples from different cities consistently overlapped (particularly from Tucson and Dubai), suggesting that while there is diversity within the household, the actual taxonomic composition of the household microbiome have some similarities, at least between these two locations. Additionally, at the species level, the taxa included in the core microbiome for each household location in Dubai and Tucson were more aligned with each other and contained skin-associated bacteria, whereas those in Mysuru were more diverse, with much higher levels of enteric bacteria in the core. Despite Tucson and Dubai being geographically separated by 8,370 miles across the globe, the households within the cities maintained similar traits in the microbiome. One hypothesis for this could be the climate of the locations – Dubai and Tucson both have hot, dry climates, classified as arid, desert, and hot (BWh) by the Köppen-Geiger climate system, whereas the climate in Mysuru is considered arid, steppe, and hot (BSh)^45^. Climate was shown to strongly drive the indoor microbiome composition in a study by Wang et al.^46^, so this is a possible explanation for the differences in bacterial diversity between the two climate types. Additionally, during sampling, many households in Mysuru had the windows open, whereas in Tucson and Dubai they were closed, reducing environmental microbes from entering, which might also be due to the climates of the regions. It has been observed that lower socioeconomic status leads to a more diverse gut microbiome^47^. Given the strong impacts which the human microbiome has on shaping the built microbiome, socioeconomic status could also play a subtle role in shaping the built microbiome. This could also be driven by other factors like how much time can be dedicated to cleaning or how many cleaning products can be purchased and used. Therefore, other possibilities could include differences in the level of industrialization in the three cities, where Dubai and Tucson are more developed, differences in cleaning products or habits, or cultural habits between the locations. However additional studies are needed to completely decipher the drivers associated with variation in the household microbiomes between the three cities. A high level of diversity across the globe has been observed in the built microbiome, and the drivers behind this diversity should be investigated further, as previous research has suggested that cultural practices, socioeconomic status, climate, and hygiene habits have shown to play a large role in shaping the built microbiome^46^.

Overall, the taxa which composed the microbiome of different household locations were diverse, but patterns could be found in locations with similar uses or had similar frequencies of water contact or cleaning. Our study found that the microbiome within the household was significantly variable across the different household locations regardless of the country. Alpha diversity varied throughout the house in each city but was found to typically be highest under the toilet rim and on the kitchen and bathroom sinks. A potential driver behind this could be moisture levels - previous studies found that surface material and moisture levels influenced the diversity of microbes and even the presence and survivability of pathogens in a specific location^3, 14, 48, 49^. Lax et al found that repeated wetting of a surface lead to reduced microbial diversity^50^ - this is reflected in our data through the toilet bowl and showerhead locations, where water routinely passes over the smooth ceramic or metal surface but does not pool, allowing the surface to dry and decreasing survivability for less ‘hardy’ microbes as they are constantly rinsed off. Another potential driver of diversity between the household locations would be when they were last cleaned, used, or touched by an individual living in the house, as external stimuli may momentarily impact the microbiome, even if it normally maintains a stable composition^37^. This is difficult to measure in our study as we did not collect this data, and touch patterns on household surfaces are incredibly variable. A few taxa were found universally across the household - for example, many samples included taxa in the family *Moraxellaceae*, which was found to be part of the core microbiome in many locations in this study, and has also been identified in other research studies looking at different household samples such as kitchen sponges and washing machines^14, 45, 48^. More research should be done to investigate how cleaning habits and location niches as well as abiotic factors like climate impact the microbiome across the built environment, as this is likely a driver of the microbiome composition worldwide.

One limitation of this study was that a few of the household locations sampled in Tucson and Mysuru were not sampled in Dubai households due to access, meaning that exact comparisons could not be made across all ten household locations in the three cities. Addressing this inconsistency would provide a clearer understanding of what taxa are found in different household locations across the globe. Despite this, most of the locations were collected consistently across the 30 houses, so this issue only impacted a few sites. Additionally, due to differences in the capacities of the lab in Tucson versus Mysuru, different sequencing kits were used, which may impact the depth and breadth of the sequenced data. To address this, most samples were sequenced multiple times to get a standard number of reads per sample across the dataset, however this does not fully reduce the bias between different sequencing library preparations. Analysis focused on relative abundance of taxa, which may mis-represent the true abundance of certain taxa within each sample. This study did not analyze cleaning, cultural habits, or other factors like flooring type or presence of pets within the houses. Information such as how often the counters and bathroom were cleaned, whether windows were kept open or closed, if pets are tracking in dirt from outside, and more could add valuable context to the microbiome data. This could potentially explain why certain taxa were present in some locations but not others and explain some of the influence of human activity on bacterial diversity. We only sampled houses with western-style toilets (ceramic sitting toilets, as opposed to squat toilets) and indoor kitchens, two features which may not be common globally and may harbor a very different bacterial profile. The samples were collected in different months of the year (winter in Tucson and Mysore, and early fall in Dubai) so seasonal differences could impact the microbiome. Finally, two of the three study locations had very similar climates (Tucson and Dubai), so sampling another location with a different climate could yield data which encompassed the household microbiome more wholly across the globe. To improve this study, more information should be collected on household habits to identify any influential lifestyle variables on the microbiome, the sample collection should be done at the same time of year, and additional study locations with varying climate could be added to increase the scope of the analysis. Finally, further studies could look at the functional capabilities of the microbiome and could assess changes in the microbiome over time, in contrast to our cross-sectional sampling.

The indoor household microbiome is dynamic and diverse, and the taxa exploit this niche are determined both by surface use and type and geographic location (cultural and environmental factors). It can also be largely influenced by the presence of humans in the house, with implications leading to the most abundant phyla in the built microbiome being human-associated^16, 40^. Similar to previous studies on household microbiome in various countries the same phyla were found in Tucson, Dubai, and Mysuru, suggesting that the main phyla in the household microbiome are human-associated (Actinomycetota, Bacillota, Gammaproteobacteria, and Pseudomonadota) across the majority of large cities^1, 3, 7, 11, 13, 14^. The microbiome of different locations throughout the house was more variable, yet similarities were seen between the microbes found across the house in Dubai and Tucson, which have similar climates, as well as similar taxa were found in household locations which experience the same moisture conditions or in locations that were adjacent to each other. Likely, a great number of external factors are driving the diversity in the household, such as climate, cleaning regimes, and different cultural practices. Future studies focused on the influence of human habits or control over some aspects which could potentially influence the microbiomes, like the presence of air conditioning in the household, is necessary to decipher what factors are driving the composition of the microbiome, aside from simply the presence of humans in the house. Overall, our study found that the household bacterial microbiome in Tucson, Mysuru, and Dubai were composed of human-associated phyla, similar to what was found in other cities during previous studies, but the locations within houses in each city (and the cities themselves) had more distinct bacterial communities at the family and genus level, which may be selected by the niche of the specific location in the house. The identification of pathogens (e.g., *Escherichia coli, Salmonella enterica, Shigella flexneri, Pseudomonas aeruginosa, Francisella halioticida, Moraxella osloensis*, and *Staphylococcus epidermidis)* within household niches highlights the importance of evaluating their ecological behavior within the broader built microbiome. Future research should investigate how targeted, surface-specific hygiene interventions (bi-directional hygiene or ‘bygiene’ approaches) influence the fate, survivability, and ecological balance of these pathogens to mitigate risk while preserving a healthy indoor microbiome^51^. While similarities can be drawn between household locations in one city and another, more research must be conducted to determine the specific factors, such as cultural practices, surface types, climate, and cleaning and hygiene habits that drive the composition of the microbiome across the house. Characterizing the microbiome of the household allows for an assessment of the risk to public health as humans are exposed to the household microbiome on a daily basis.

## Conflicts of Interest

The authors declare that there are no conflict of interest.

## Funding Information

This work was completed with funding from Reckitt Benckinser LLC.

## Acknowledgements

Thank you to the volunteers in Tucson, Dubai, and Mysuru who generously allowed for sample collection from their household.

## Notes

https://github.com/carolinescranton01/Global_Household_Microbiome

### Competing Interest Statement

This work was completed with funding from Reckitt Benckinser LLC, and authors M. Khalid Ijaz and Frank Y. Xu are employess of Reckitt Benckinser LLC.

