## Supplementary Figures for "Characterization of household microbiomes from three unique cities around the world"

**Supplemental Figure 1**

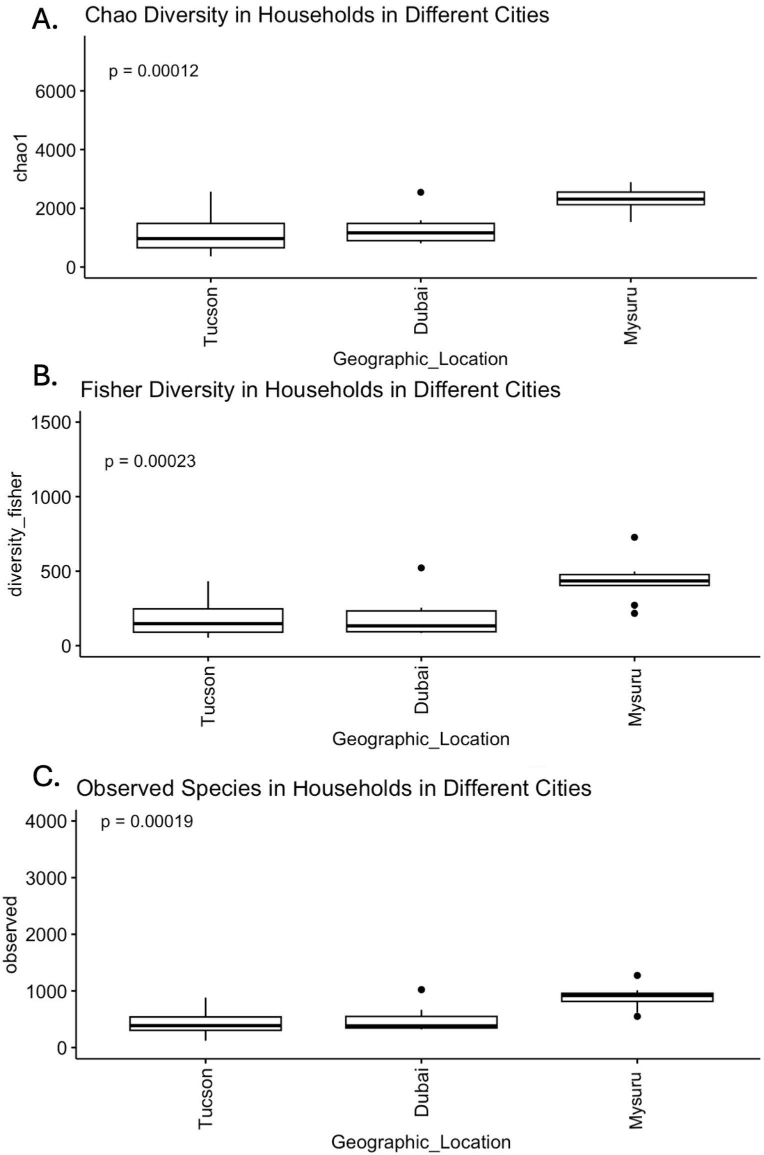


**Supplementary Figure 1. Other alpha diversity metrics between the household microbiome of the three cities overall. A: Chao diversity index, highly significant (ANOVA, p<0.01). B: Fisher diversity index, highly significant (ANOVA, p<0.01). C: Observed species index, highly significant (ANOVA, p<0.01).**

### Supplemental Figure 2

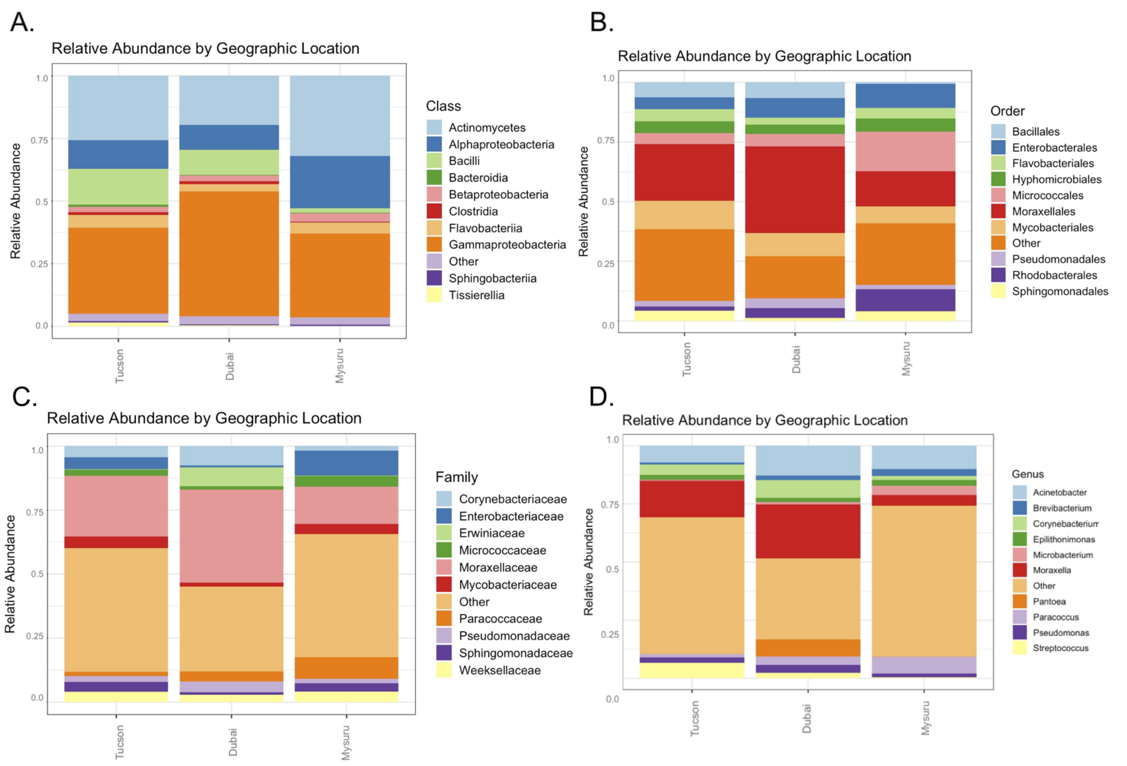


**Supplementary Figure 2. The taxonomic makeup between the household microbiome of the three cities overall at different taxonomic levels. The relative abundance of the top 10 most abundant taxa is shown. A: Taxonomic level = Class. B: Taxonomic level = Order. C: Taxonomic level = Family.**

**Supplemental Figure 3**

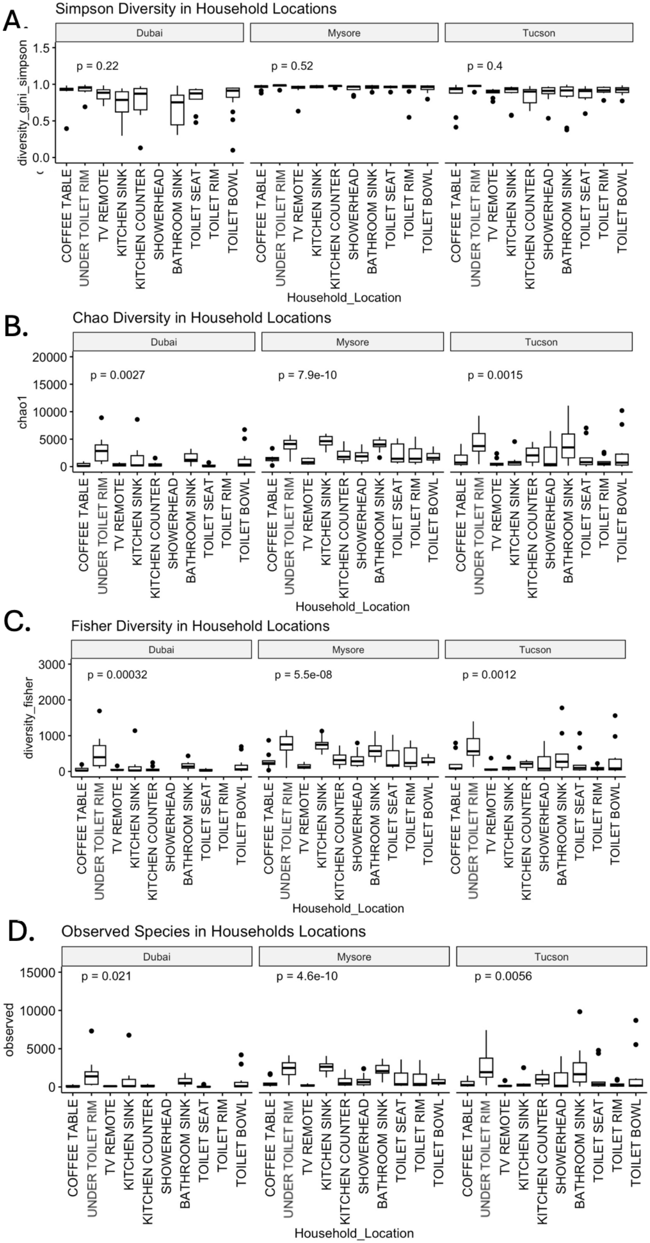


**Supplementary Figure 3. Other alpha diversity metrics between the household microbiome of the three cities overall. A: Simpson diversity index, not significant in any city. B: Chao diversity index, highly significant in all cities (ANOVA, p<0.01). C: Fisher diversity index, highly significant in all cities (ANOVA, p<0.01) One value excluded – Dubai House 7 toilet seat, value = 1.07E9. D: Observed species index, significant in Dubai (ANOVA, p<0.05), highly significant in Mysuru and Tucson (ANOVA, p<0.01).**

### Supplemental Figure 4

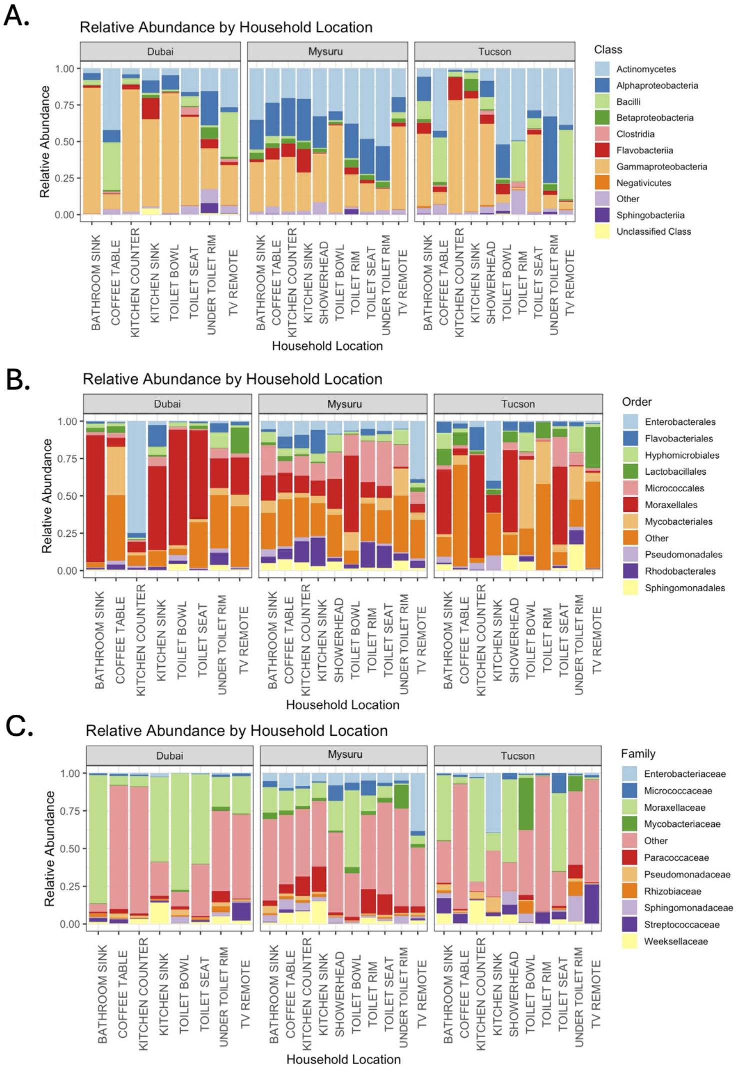


**Supplementary Figure 4. The taxonomic makeup at each household location within the three cities at different taxonomic levels. The relative abundance of the top 10 most abundant taxa is shown. A: Taxonomic level = Class. B: Taxonomic level = Order. C: Taxonomic level = Family.**
